# Motor neurons involved in fine motor control are labeled by tracing *Atoh1*-lineage neurons in the spinal cord

**DOI:** 10.1101/2020.03.20.000588

**Authors:** Osita W. Ogujiofor, Iliodora V. Pop, Felipe Espinosa, Razaq O. Durodoye, Michael L. Viacheslavov, Rachel Jarvis, Mark A. Landy, Gordon Fishell, Robert P. Machold, Helen C. Lai

## Abstract

Motor neurons (MNs) innervating the digit muscles of the intrinsic hand and foot (IH and IF) control fine motor movements. Previous studies suggest that the IH and IF MN pools have a unique developmental history in comparison to limb MN pools. Consistent with having this unique development, we find that the IH and IF MN pools are labeled postnatally using a CRE knock-in mouse line of *Atoh1*, a developmentally expressed basic helix-loop-helix (bHLH) transcription factor, while limb-innervating MN pools are not. Approximately 60% of the IH and IF MN pools are labeled and are a mixture of alpha and gamma-MNs. In addition, because *Atoh1* is known developmentally to specify many cerebellar-projecting neurons, we tested the hypothesis that IH and IF MNs can send axon collaterals to the cerebellum as a mechanism of corollary discharge. Using intersectional genetic, viral labeling, and retrograde labeling strategies, we were unable to provide evidence in support of this idea. As a secondary finding of our viral labeling experiments, we report here that injection of both AAV and Lentiviruses in the periphery can cross the blood-brain barrier to infect Purkinje cells within the central nervous system. Altogether, though, we find that labeling of the IH and IF motor neurons using the *Atoh1* CRE knock-in mouse suggests that IH and IF MNs have a unique developmental history and that this mouse strain might be a useful tool to target these specific sets of neurons allowing for functional studies of fine motor control.

**Significance Statement:** Motor neurons (MNs) of the intrinsic hand and foot (IH and IF) are labeled postnatally using a CRE knock-in mouse line of the basic helix-loop-helix (bHLH) transcription factor *Atoh1* indicating a unique developmental history. We tested whether IH and IF MNs send axon collaterals rostrally to the cerebellum as a mechanism of direct corollary discharge from MNs, but the question remains unresolved. As a resource for the community, we report that injection of both AAV and Lentiviruses in the periphery can cross the blood-brain barrier and infect Purkinje cells within the central nervous system.

## Introduction

Fine motor skills, such as writing or sewing, require exquisite control of the motor neurons (MNs) innervating the digits of the hand. Over vertebrate evolution, unique molecular pathways are involved in the elaboration of digits (Shubin et al., 1997). Correspondingly, the MNs innervating the distal digits have unique developmental programs compared to the neighboring MNs of the lateral motor column (LMC) that innervate limb muscles (Mendelsohn et al., 2017). Precisely how digit-innervating MNs adopt their unique identities is unclear. Insight into the development and connectivity of digit-innervating MNs could reveal distinct functions of these MNs in fine motor behavior.

Developmentally, all MNs derive from a progenitor domain expressing the basic helix-loop-helix (bHLH) oligodendrocyte transcription factor 2 (*Olig2*) in the ventral neural tube (Lu et al., 2002). Unexpectedly, we found that the digit-innervating MNs were labeled using CRE-loxP lineage tracing of the bHLH transcription factor atonal homolog 1, *Atoh1*, a transiently expressed gene in the dorsal-most part of the developing neural tube that specifies spinal cord neurons that project rostrally to the hindbrain (Bermingham et al., 2001; Gowan et al., 2001; Sakai et al., 2012; Yuengert et al., 2015). Here, we explored the features of the digit-innervating MNs labeled by *Atoh1* CRE-LoxP lineage tracing.

In the motor field, corollary discharge is a mechanism by which copies of descending motor signals are sent back to the proprioceptive sensory pathways to distinguish self-generated movements from externally generated ones (Sperry, 1950; von Holst and Mittelstaedt, 1950; Crapse and Sommer, 2008). In mammals, to the best of our knowledge, there has been no evidence of corollary discharge occurring at the level of MNs themselves, although retrograde labeling studies of spinocerebellar neurons have reported cerebellar-projecting neurons whose cell bodies reside in the MN lamina IX of the spinal cord (Matsushita and Hosoya, 1979; Matsushita et al., 1979; Terman et al., 1998). In addition, Cooper and Sherrington in their initial description of ascending projections from the spinal cord observed “large cells” in the ventral horn that degenerated upon cutting their ascending axons (Cooper and Sherrington, 1940).

The fact that *Atoh1*-lineage neurons include many cerebellar-projecting neurons such as those of the spinocerebellar system, pedunculopontine tegmentum, pontine nuclei, lateral reticular nucleus, and external cuneate nucleus (Rose et al., 2009), raised the intriguing possibility that *Atoh1* CRE-LoxP lineage traced MNs might indicate a function of ATOH1 in these MNs to specify axon collaterals to the cerebellum. Evidence of axon collaterals to the cerebellum from MNs, specifically the MNs involved in fine motor control, would support a model of corollary discharge directly from MNs themselves. We attempted to test this model using intersectional genetic, viral tracing, and retrograde tracing strategies, but each approach had its caveats. Here, we characterize the *Atoh1* CRE-LoxP lineage traced MN populations and report our efforts testing a model of corollary discharge from fine motor control MNs.

## Materials & Methods

### Mouse strains

The following mouse strains were used: *Atoh1*^*Cre/+*^ knock-in (Yang et al., 2010), *R26*^*LSL-tdTom/+*^ (Ai14)(Madisen et al., 2010), *R26*^*LSL-FSF-tdTom/+*^ (Ai65)(Madisen et al., 2015), *Hoxa4::Cre* (Huang et al., 2012), *Chat*^*IRES-Cre*^ (Rossi et al., 2011), *Cdx2::FLPo* (Bourane et al., 2015). The *Chat*^*IRES-FLPo*^ mouse was generated by Gord Fishell and Rob Machold (unpublished). Briefly, the IRES-FLPo-pA cassette was knocked into the 3’UTR immediately following the *ChAT* stop codon in B4 ES cells (C57Bl/6). Following germline transmission of the correctly targeted ChAT-IRES-FLPo allele, the Neo cassette (LoxP flanked) was removed by crossing the mice with the CMV-Cre deleter line (JAX # 006054) prior to use. All mice were outbred and thus, are mixed strains (at least C57Bl/6J and ICR). *Atoh1*^*Cre/+*^ knock-in mice crossed to reporter mice were screened for “dysregulated” expression as previously reported (Yuengert et al., 2015). All animal experiments were approved by the Institutional Animal Care and Use Committee at UT Southwestern.

### Tissue processing

Embryos were timed as E0.5 on the day the vaginal plug was detected and P0 on the day of birth. Pregnant females were euthanized with CO_2_ and cervical dislocation, embryos dissected out of the uterus, and spinal cords dissected out. Embryonic spinal cords (E14.5) were fixed in 4% paraformaldehyde (PFA)/PBS for 2-3 hrs at 4°C. Early postnatal animals (younger than P7) were cooled on ice, decapitated, their spinal cords dissected out, and fixed in 4% PFA/PBS for 2 hours at 4°C. Mice older than P14 were anesthetized with Avertin (2,2,2-Tribromoethanol) (0.025-0.030 mL of 0.04 M Avertin in 2-methyl-2-butanol and distilled water/g mouse) and transcardially perfused, first with 0.012% w/v Heparin/PBS and then 4% PFA/PBS. A dorsal or ventral laminectomy exposed the spinal cord to the fixative. The spinal cords were fixed for 2 hrs and the brains overnight at 4°C. Tissue was washed in PBS for at least one day and cryoprotected in 30% sucrose dissolved in deionized water. Tissue was marked with 1% Alcian Blue in 3% acetic acid on one side to keep orientation and were then embedded in OCT (Tissue-Tek Optimal Cutting Temperature compound). Tissue was sectioned using a Leica CM1950 Cryostat.

### Immunohistochemistry and confocal imaging

Cryosections (30-40 μm) were blocked with PBS/1-3% normal goat or donkey serum/0.3% Triton X-100 (Sigma) for up to 1 hour at room temperature (RT) and incubated overnight with primary antibody at 4°C. After washing 3 times with PBS, the appropriate secondary antibody (Alexa 488, 567, and/or 647, Invitrogen) was incubated for an hour at RT. Sections were rinsed 3 times in PBS, mounted with Aquapolymount (Polysciences Inc.), and coverslipped (Fisher). The following primary antibodies and dilutions were used: 1:500 rabbit anti-dsRed (Clontech), 1:100 goat anti-CHAT (Millipore Sigma), 1:1000 rabbit anti-MMP9 (Abcam), 1:8000 rabbit anti-HB9 (gift of Dr. Sam Pfaff, Salk Institute), 1:3000 guinea pig anti-CPNE4 and 1:8000 guinea pig anti-FIGN (gifts of Dr. Tom Jessell, Columbia Univ.), 1:500 mouse anti-NEUN (Millipore Sigma), 1:100 mouse anti-ERR3 (R&D Systems), 1:3000 alpha-bungarotoxin 488 (Invitrogen), 1:1000 rabbit anti-Syntaxin1 (gift of Thomas Südhof, Stanford University), 1:1000 guinea pig anti-VGLUT1 (Millipore Sigma), 1:1000 guinea pig anti-VGLUT2 (Millipore Sigma), 1:1000 goat anti-VACHT (Millipore Sigma). Sections were referenced to the Mouse Brain Atlas (Paxinos and Franklin, 2007) and Christopher Reeves Spinal Cord Atlas (Watson et al., 2009).

Fluorescent images were taken on a Zeiss LSM710 or LSM880 confocal microscope with an appropriate optical slice (0.5-10 μm) depending on the image. Images were pseudocolored using a magenta/green/blue or magenta/yellow/cyan color scheme using Adobe Photoshop (Adobe) or Fiji (Schindelin et al., 2012).

### CTB muscle injections

Mice aged P14 were anesthetized using isoflurane and prepared for injections into muscle. An approximate total of 500-750 nL of Cholera toxin subunit B (CTB) AlexaFluor 488 or 647 Conjugate (Invitrogen) (Nanoject II, Drummond Scientific) was injected into 2-3 different locations in the left forepaw (Intrinsic Hand (IH) MN pool) or hindpaw (Intrinsic Foot (IF) MN pool), or 3-4 different locations for the gastrocnemius (GS) or tibialis anterior (TA) in 50.6 nL increments. Spinal cords were harvested 5 days after injection.

### Viral Injections

Mice aged P4-5 or P14-15 were anesthetized using isoflurane (Henry Schein) and prepared for injections into the hindpaw. The hindpaw was shaved if needed and 70% ethanol and betadine (Avrio Health L.P.) applied. For P4-5 pups, virus was injected in a single location through the skin. For P14-15, a midline incision was made on the dorsal surface of the hindpaw. A total of 200-250 nL of AAV8-hSyn-GFP-Cre (UNC Vector Core, 6.5 x 10^12^ Vg/mL) or Lenti^FugE^-Cre, a pseudotyped lentivirus mediating the expression of CRE, was injected in 50.6 nL increments (Nanoject II, Drummond Scientific) with 1-2 min between injections at 2-3 different locations in the left hindpaw for P14-P15 animal and at a single location through the skin in P4-5 pups. Lenti^FugE^-Cre was pseudotyped with a fusion glycoprotein enabling efficient retrograde axonal transport (Kato et al., 2014). To generate Lenti^FugE^-Cre, *Cre* was sub-cloned into the third generation HIV-based lentivirus vector under the control of a synapsin promoter (FSW-*Cre*). FSW-*Cre* was co-transfected into HEK293 cells with three packing plasmids, pMDLg/pRRE, pRSV-Rev and pCAGGS-FuG-E to generate Lenti^FugE^-Cre, which was concentrated with ultracentrifugation to 2.0 x 10^12^ Vg/mL. The incision was closed with surgical glue (Henry Schein). Carprofen (5 mg/kg) was administered daily 3 days after surgery. Spinal cords were harvested 21-27 days after injection.

### Fluorogold Injections

Two P32 *Atoh1*^*Cre/+*^ TOM^+^ female mice were injected with 4% (w/v) FG solution in saline (Fluorochrome). Mice were anesthetized with isoflurane and the area above and around the cerebellar region was prepared for surgery. A midline incision of 0.75 cm and a craniectomy of approximately 1 mm wide by 1.5 mm long was performed. Bilateral injections at six sites were done at (from Bregma): rostrocaudal -5.5, -5.9, and -6.3 mm and at mediolateral ± 0.2-0.4 mm. At each site, several injections in 50.6 nL increments were performed every 300 μm along the dorsoventral axis starting at -1.7 mm deep for a total of 720 nL of FG on each side in mouse #1, and 270 nL on each side in mouse #2. Animals were harvested 7 days after injection.

### Whole tissue imaging

Mouse brainstem and spinal cords were processed following the SHIELD protocol (Park et al., 2018). Tissues were cleared with SmartClear II Pro (LifeCanvas Technologies, Cambridge, MA) for several days, mounted in a gel of 0.9% agarose in EasyIndex (LifeCanvas Technologies), and then incubated in EasyIndex for refractive index matching. Tissues were imaged at 3.6X using a SmartSPIM light sheet microscope (LifeCanvas Technologies). The *Chat*^*Hoxa4*^ hindbrain sample (Movie 1) was imaged at 1.8 μm x 1.8 μm x 4 μm resolution. The *Chat*^*Hoxa4*^ spinal cord samples (Movies 2 and 3) were imaged at 1.8 μm x 1.8 μm x 2 μm resolution. The hindbrain and spinal cord samples were cut to less than 2.2 cm to fit in the imaging chamber. The spinal cord samples were cut into two pieces (cervical-thoracic and thoracic-lumbar) and cleared separately. Movies were made in arivis Vision4D 2.12.6.

### Experimental Design and Statistical Tests

For the percentage of CHAT^+^ neurons in the IH and IF MN pools that were *Atoh1*-lineage TOM^+^ neurons (Fig. 1D) and the estimated total number of CHAT^+^ neurons in the IH and IF MN pools, IH data were counted from 3-4 sections per animal from n=2 female (F) mice and IF data were counted from 3-4 sections per animal from n=3 F mice from two different litters. For estimating the total number of CHAT^+^ neurons, counts from 4 MN pools left and right side were counted from sections that represented a tenth of the MN pool. Therefore, final estimates of the total number of CHAT^+^ neurons were the final counts multiplied by ten. For the percentage of *Atoh1*^*Cre/+*^ TOM^+^ MNs that were MMP9^+^ fast twitch MNs (Fig. 1F), IH and IF data were counted from 3 sections per animal from n=2 (1 F, 1 male (M)) mice of the same litter. For the percentage of HB9^+^ neurons that were *Atoh1*-lineage TOM^+^ neurons over developmental time (Fig. 3B), IH and IF data were counted from 2-4 sections per animal from n=2 mice (from the same litter for P3 (gender not noted), P7 (2 F), and P15 (1 F, 1 M); two different litters for P23 (IH MN pool was from 1 unknown gender and 1 M, IF MN pool was from 1 unknown gender and 1 F). For the percentage of *Atoh1*-lineage TOM^+^ neurons that are ERR3^+^ or NEUN^+^ (Fig. 4C), IH and IF data were counted from 3 sections per animal from n=2 mice (1 F, 1 M) from the same litter. For *Chat*^*Hoxa4*^ MNs, 4 sections per animal were counted from n=2 mice (2 M) from two different litters. For the *Chat*^*Cdx2*^ MNs, 4 sections were counted from one female mouse. The *Chat*^*Hoxa4*^ mouse processed with SHIELD by LifeCanvas Technologies was male.

**Figure 1.**
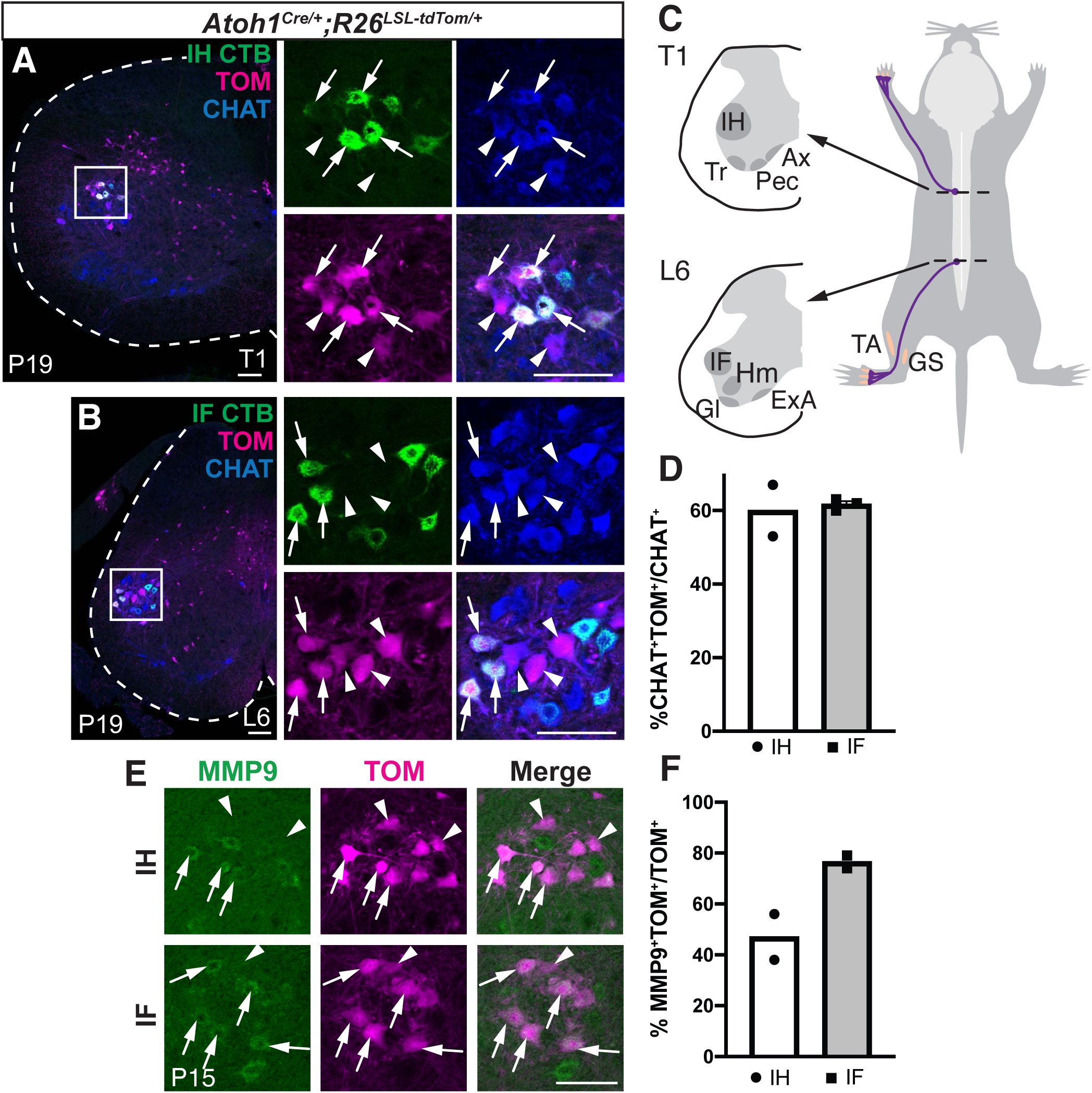
The *Atoh1*^*Cre/+*^ knock-in mouse line labels the intrinsic hand (IH) and foot (IF) motor neuron (MN) pools. (A, B) Injection of the retrograde tracer CTB-488 into the forepaw and hindpaw labels the IH and IF MN pools, which are labeled with tdTomato (TOM) when the TOM reporter mouse is crossed to the *Atoh1*^*Cre/+*^ knock-in mouse. Arrows, CTB^+^ CHAT^+^ TOM^+^; arrowheads, CTB^-^ CHAT^+^ TOM^+^. (C) Diagram of motor neuron pools at T1 and L6. (D) Quantitation of the percentage of the IH or IF MN pools that are labeled TOM^+^ in *Atoh1*^*Cre/+*^ knock-in mice. (E-F) Some of the TOM^+^ IH MNs and IF MNs are fast twitch MNs (MMP9^+^). MMP9^+^TOM^+^ arrows; MMP9^-^TOM^+^ arrowheads. See text for values in D, F. Christopher Reeve Atlas referenced for spinal cord MN pools (Watson et al., 2009). Abbrev: P, postnatal; C, cervical; T, thoracic; L, lumbar; Tr, tricep; Pec, pectoral; Ax, axial; Hm, hamstring; Gl, gluteus; ExA, external anal sphincter; TA, tibialis anterior; GS, gastrocnemius. Scale bars: 100 μm.

**Figure 2.**
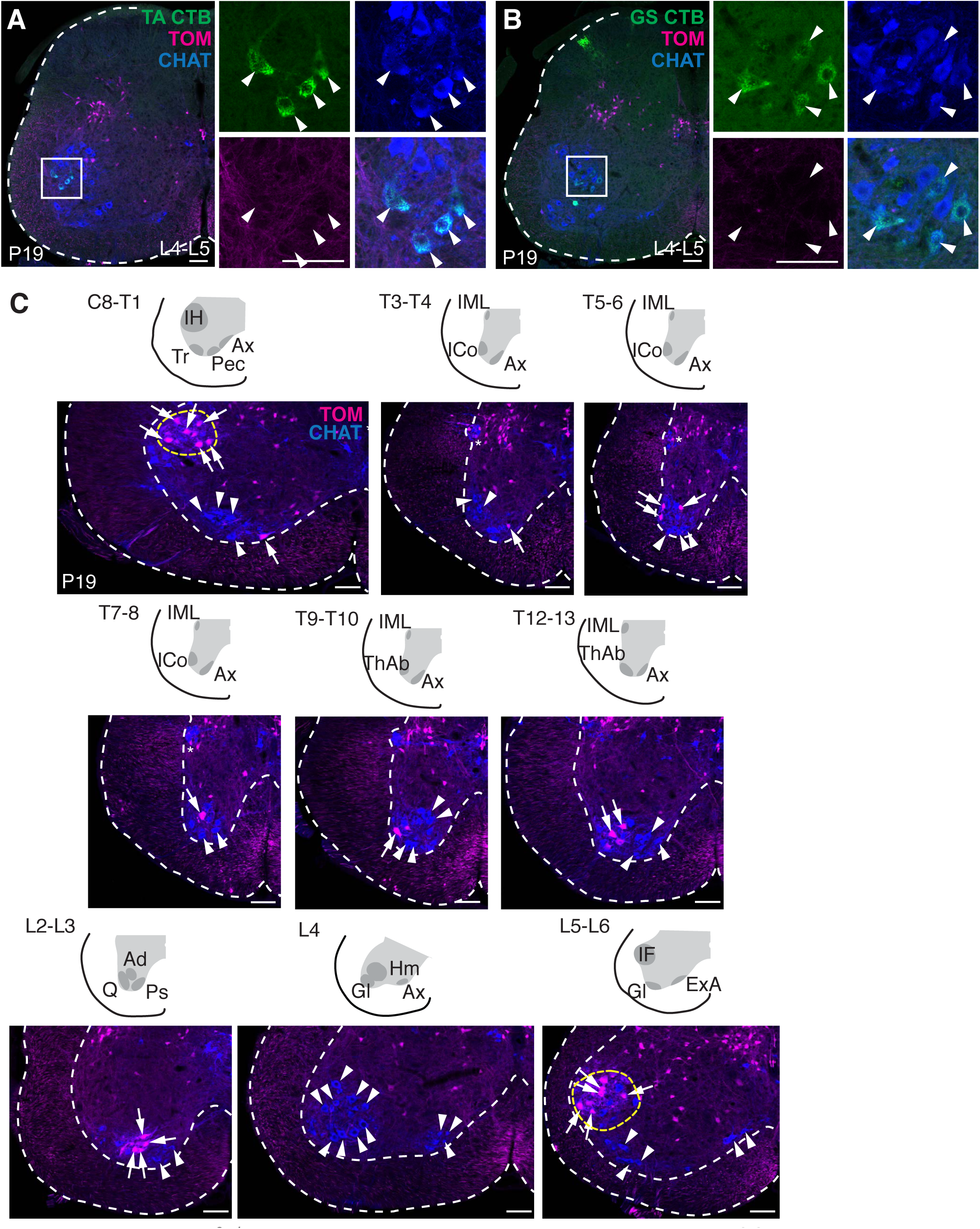
The *Atoh1*^*Cre/+*^ knock-in mouse line does not label the TA and GS MN pools and sparsely labels other MN pools. (A, B) Injection of CTB-488 into the TA and GS shows no labeling (TOM^-^, arrowheads) of these MN pools in the *Atoh1*^*Cre/+*^ knock-in mouse. (C) Representative images throughout the rostral-caudal axis of *Atoh1*^*Cre/+*^ knock-in mice crossed to the TOM reporter mouse show that TOM labels MNs mainly in IH and IF (yellow dashed lines) with sparser labeling of MNs in other MN pools (arrows). Some MN pools have no TOM^+^ expression (arrowheads). Christopher Reeve Atlas referenced for spinal cord MN pools (Watson et al., 2009). Abbrev: IML, intermediolateral nucleus; ICo, intercostal; ThAb, thoracic abductor; Q, quadricep; Ad, adductor; Ps, psoas. Scale bars: 100 μm.

**Figure 3.**
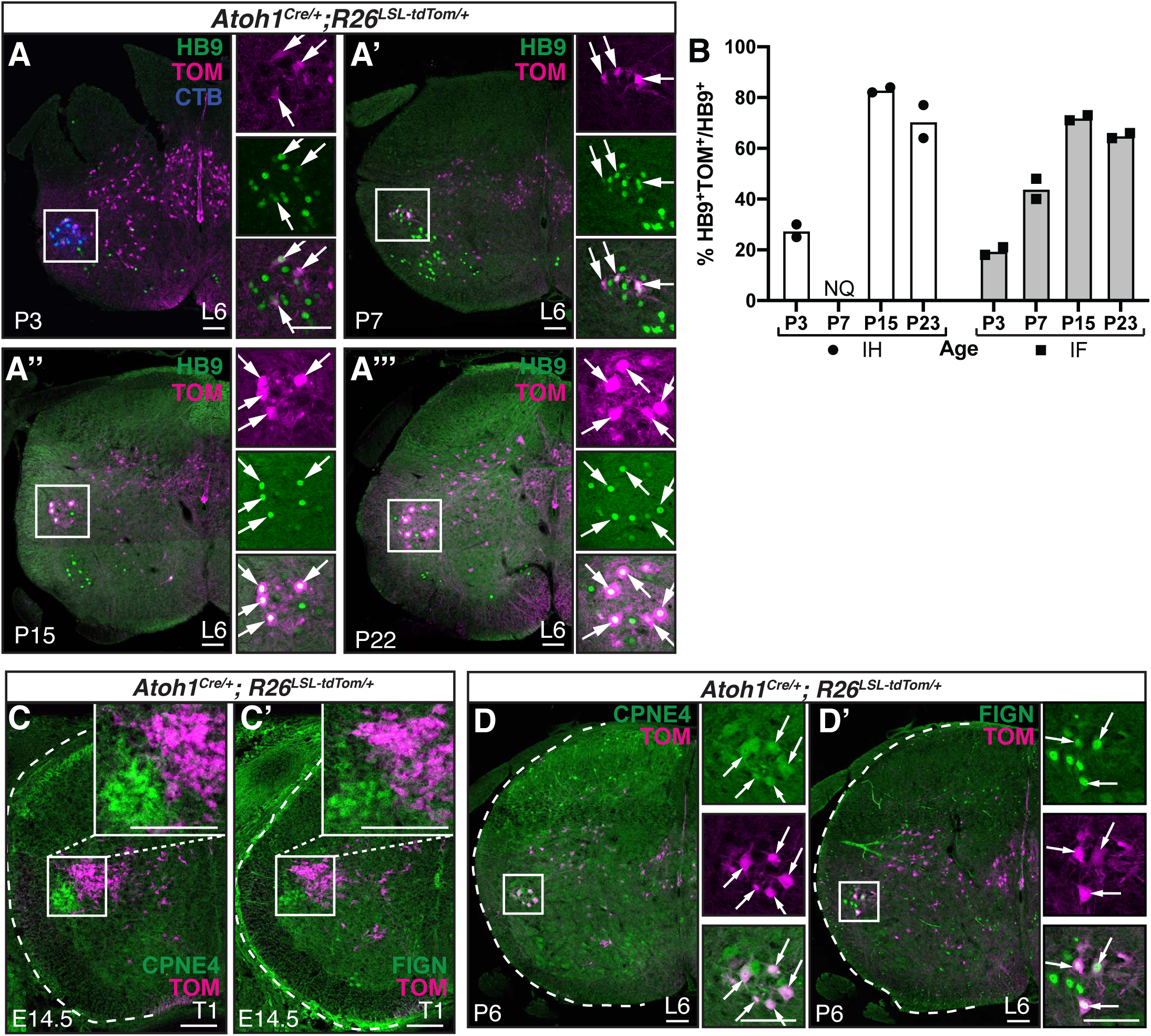
The *Atoh1*^*Cre/+*^ knock-in mouse labels IH and IF postnatally. (A-A’’’) TOM^+^ labeling of the IF MN pool at several postnatal time points. HB9^+^TOM^+^ neurons, arrows. In A, CTB (blue) was injected into the hindpaw to identify the IF MN pool. (B) Quantitation of percentage of TOM^+^ neurons in the IH or IF MN pools at several time points. (C-C’) At E14.5 (embryonic day 14.5), TOM^+^ neurons are CPNE4^-^ and FIGN^-^. (D-D’) At P6, TOM^+^ neurons are CPNE4^+^ and FIGN^+^ (arrows). Scale bars: 100 μm.

**Figure 4.**
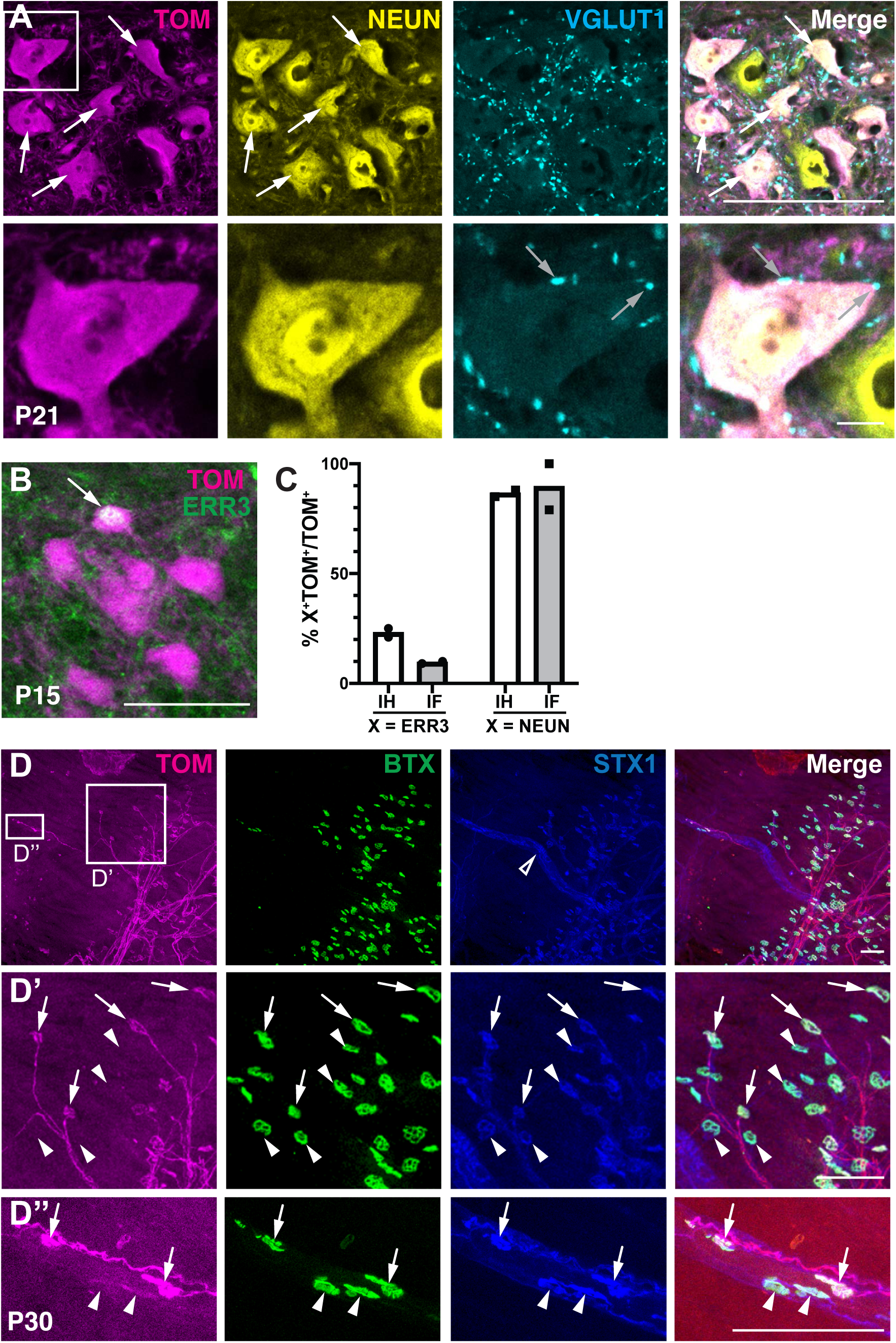
Both α- and γ-MNs are labeled in the *Atoh1*^*Cre/+*^ knock-in mouse. (A) TOM^+^ MNs in the IF MN pool are NEUN^+^ (arrows) and have closely apposed VGLUT1^+^ boutons (grey arrows). (B) Some TOM^+^ IF MNs are also ERR3^+^ (arrow). (C) Percentage of the TOM^+^ MNs in the IH and IF that are ERR3^+^ (γ-MN marker) or NEUN^+^ (α-MN marker). (D-D’’) TOM^+^ axons in the hindpaw lumbrical muscle show the neuromuscular junction innervating extrafusal muscle (D’, arrows, BTX^+^STX1^+^TOM^+^). TOM^+^ axons also innervate the intrafusal muscle spindle (D, open arrowhead; D’’, arrows, BTX^+^STX1^+^TOM^+^). Arrowheads indicate motor endplates that are TOM^-^. Scale bars: 100 μm, inset in A is 10 μm.

No statistical tests were required as quantitation of the percentage of particular markers in any given MN pool were not directly compared to each other. For samples with n=2, the mean is shown with no standard error of the mean (SEM) since the range between the two data points equals the mean ± SEM.

## Results

### Atoh1^Cre/+^ knock-in mice label MN pools involved in fine motor control

We observed using CRE-lineage tracing strategies (*Atoh1*^*Cre/+*^ knock-in mice (Yang et al., 2010) crossed to tdTomato (TOM) reporter mice (*R26*^*LSL-tdTom*^, Ai14)(Madisen et al., 2010)) that subsets of motor neurons (MNs) expressing choline acetyl transferase (CHAT) were labeled in the spinal cord (Fig. 1A, B; both arrows and arrowheads, CHAT^+^TOM^+^). Based on the anatomical location of the MN pools along the rostral-caudal axis labeled in *Atoh1*^*Cre/+*^ mice, we predicted that the *Atoh1*^*Cre/+*^ line labeled MNs of the intrinsic hand (IH) and foot (IF) in thoracic 1 (T1) and lumbar 6 (L6) areas of the spinal cord (Fig. 1C)(Watson et al., 2009). We tested our prediction by injecting the forepaw and hindpaw with the retrograde tracer choleratoxin B conjugated to Alexa 488 (CTB-488), which labeled the IH and IF MN pools. We found that the *Atoh1*^*Cre/+*^ TOM^+^ MNs indeed labeled the IH and IF MN pools (Fig. 1A, B, arrows) and made up 60.0% (mean, range 53-67%, n=2) and 61.7% ± 0.9% SEM (mean, n=3) of the IH and IF MN pools, respectively (Fig. 1D)(See Experimental Design and Statistical Tests section of the Materials and Methods for details of quantitation throughout the article). We estimate that the total number of CHAT^+^ MNs at P19 in the IH and IF MN pools on one side is IH 370 ± 63 SEM neurons (counts from 4 MN pools left and right side from n=2 mice) and IF 335 ± 5 SEM neurons (counts from 4 MN pools left and/or right side from n=3 mice). In addition, we found that a subset of the TOM^+^ MNs were fast twitch MNs (Fig. 1E-F, arrows, MMP9^+^TOM^+^; IH 47.0% (mean, range 38-56%, n=2) and IF 76.5% (mean, range 74-79%, n=2))(Kaplan et al., 2014). Note that the other TOM^+^ cell bodies in the intermediate spinal cord are from other *Atoh1*-lineage interneurons involved in the proprioceptive system (Yuengert et al., 2015).

To see whether the labeling of *Atoh1*^*Cre/+*^ TOM^+^ MNs was specific to the IH and IF MN pools, we injected CTB-488 into the tibialis anterior (TA) and gastrocnemius (GS) muscles and found that those MN pools did not have any TOM^+^ MNs (Fig. 2A, B, arrowheads). In addition, we sampled sections throughout the rostral-caudal axis of the spinal cord in *Atoh1*^*Cre/+*^ mice. We found that other MN pools had TOM^+^ expression (Fig. 2C, arrows, CHAT^+^TOM^+^). However, the TOM^+^ MN labeling was enriched in the IH and IF MN pools (Fig. 2C, C8-T1 and L5-6, yellow dashed lines).

### Atoh1^Cre/+^ knock-in mice label IH and IF MN pools postnatally

Given that MNs are derived from an *Olig2*-expressing progenitor domain in the ventral neural tube (Lu et al., 2002; Lai et al., 2016), we tested whether the *Atoh1*^*Cre/+*^ line was labeling IH and IF MNs early in embryonic development, which would suggest co-expression with the *Olig2*-expressing domain, or if there was a previously unreported late expression of *Atoh1* in the IH and IF MNs. We found that at embryonic day 14.5 (E14.5) when the IH and IF MNs first start expressing the unique markers Copine-4 (CPNE4) and Fidgetin (FIGN) (Mendelsohn et al., 2017), the IH and IF MN pools were not yet TOM^+^ (Fig. 3C, C’). In contrast, at postnatal time points, we found that colocalization of TOM^+^ MNs in the IH and IF MN pools with the homeobox transcription factor, HB9, marking MN pools, started around P3 (IH 28% (range 25-30%), IF 20% (range 18-21%), mean, n=2) and gradually increased to about 70-80% by P15 (P7: IH not quantitated (NQ), IF 44% (range 40-48%); P15: IH 83% (range 82-84%), IF 72% (range 71-73%); P23: IH 71% (range 64-77%), IF 65% (range 64-66%); mean, n=2)(Fig. 3A-A’’’, arrows, and Fig. 3B). In addition, we confirmed that the IF TOM^+^ MNs colocalized with the specific markers CPNE4 and FIGN postnatally (Fig. 3D, D’). To detect postnatal *Atoh1* expression in the IH and IF MNs, we performed *in situ* hybridization (Gowan et al., 2001 for ISH probe) and RNAscope of *Atoh1* at P14-P15 and P22, but were unable to detect any signal at the mRNA level (unpublished observations). Taken together, it is likely that the IH and IF MN pools derive from an *Olig2*-expressing progenitor domain and are labeled postnatally in the *Atoh1*^*Cre/+*^ line.

### IH and IF MN pools labeled with Atoh1^Cre/+^ knock-in mice are both α- and γ-MNs

Because *Atoh1*^*Cre/+*^ TOM^+^ MNs are enriched in only a subset (∼60%) of IH and IF MNs, we tested whether they were demarcating a specific type of MN (α or γ). α-MNs innervate the striated extrafusal muscle, are marked by the neuronal marker, NEUN, and receive vesicular glutamate transporter 1 (VGLUT1^+^) proprioceptive inputs (Friese et al., 2009; Manuel and Zytnicki, 2011; Ashrafi et al., 2012). γ-MNs innervate the intrafusal muscle spindles and express Estrogen Related Receptor gamma (ERR3^+^)(Friese et al., 2009). Immunostaining for α- and γ-MN markers, we found that *Atoh1*^*Cre/+*^ TOM^+^ MNs are a mixture of both α- and γ-MNs (Fig. 4A, B, NEUN^+^TOM^+^ and ERR3^+^TOM^+^, arrows). The NEUN^+^ *Atoh1*^*Cre/+*^ TOM^+^ MNs also received VGLUT1^+^ proprioceptive inputs (Fig. 4A). Previous reports found approximately 30% of MNs are γ-MNs and 70% are α-MNs (Friese et al., 2009). We found that the *Atoh1*^*Cre/+*^ TOM^+^ MNs were IH 23.0% (range 21-25%) and IF 9.5% (range 9-10%) γ-MNs (ERR3^+^TOM^+^/TOM^+^) and IH 86.5% (range 85-88%) and IF 89.5% (range 79-100%) α-MNs (NEUN^+^TOM^+^/TOM^+^)(Fig. 4C, mean, n=2, note that ERR3 and NEUN counts were performed on different sections). Our results suggest a slight enrichment of α-MNs in the IH and IF MN pools that may reflect endogenous differences in α- and γ-MN distribution in IH and IF MN pools or a preference for labeling α-MNs in the *Atoh1*^*Cre/+*^ mouse line. Furthermore, imaging of the hindpaw lumbrical muscle found TOM^+^ axons innervating both the extrafusal (Fig. 4D’, arrows) and intrafusal muscle (Fig. 4D’’, arrows). Bungarotoxin (BTX^+^) identifies the neuromuscular junctions (NMJs) and syntaxin (STX1^+^) identifies the muscle spindle (Fig. 4D, open arrowhead) and NMJs. Note that not all NMJs are TOM^+^ (Fig. 4D’-D’’, arrowheads) consistent with the fact that only ∼60% of the IH and IF MN pools are TOM^+^.

### Ascending projections from caudal cholinergic neurons

The *Atoh1* transcription factor is known to specify many cerebellar-projecting neurons (Rose et al., 2009). Given the labeling of the IH and IF MNs in the *Atoh1*^*Cre/+*^ mouse line, we wanted to test whether IH and IF MNs could send ascending axon collaterals, potentially to the cerebellum. To this end, we pursued two intersectional genetic strategies to label caudal cholinergic neurons (Fig. 5A, G).

**Figure 5.**
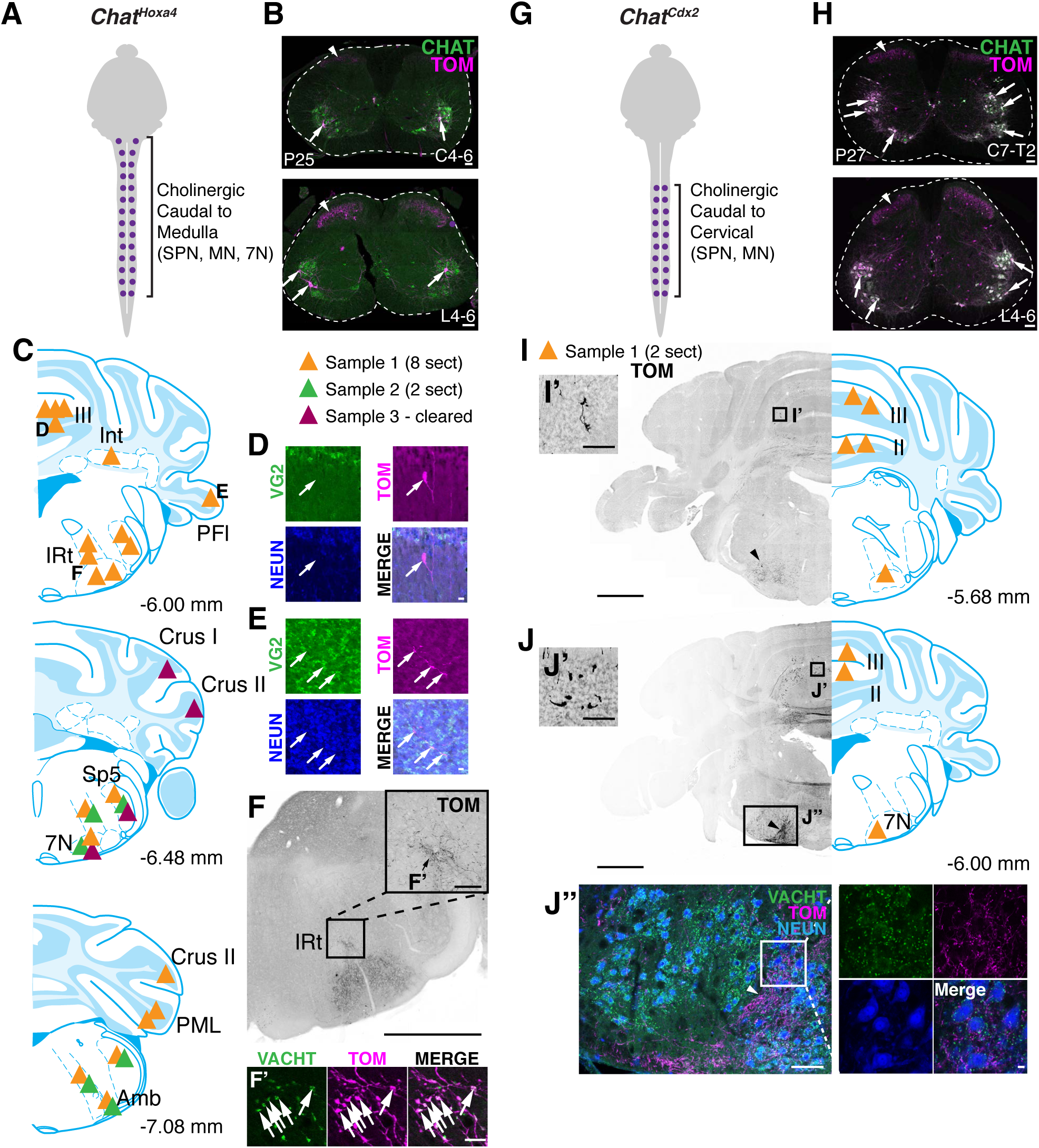
Intersectional genetic labeling of caudal cholinergic neurons finds ascending projections to the cerebellum. (A) Diagram of *Chat*^*Hoxa4*^ intersectional cross that labels cholinergic neurons caudal to the medulla. (B) Sparse TOM^+^ (arrows) labeling of MNs (CHAT^+^, green) in *Chat*^*Hoxa4*^ mice. Axonal terminations from sensory neuronal labeling are seen in the superficial dorsal horn (arrowheads). (C) Summary of data from three samples of where TOM^+^ processes are seen in the hindbrain. Number of sections (sect) analyzed is stated. (D) Representative image of TOM^+^ process (arrow) seen in vermis III that is VGLUT2 (VG2) negative. Maximum Intensity Projection (MIP) of 21 μm. (E) Representative image of TOM^+^ process (arrows) in PFl that is VG2^-^ and courses between the NEUN^+^ granule cells. Maximum Intensity Projection (MIP) of 30 μm. (F-F’) Processes in IRt express VACHT (arrows). (G) Diagram of *Chat*^*Cdx2*^ intersectional cross that labels cholinergic neurons caudal to the mid-cervical spinal cord. (H) Caudal cholinergic neurons are well-labeled in *Chat*^*Cdx2*^ mice (arrows, TOM^+^CHAT^+^). Note that axonal terminations from sensory neuronal labeling are also seen in the superficial dorsal horn (arrowheads). (I-I’, J-J’) TOM^+^ processes are seen in the folia II and III vermis of *Chat*^*Cdx2*^ mice. TOM^+^ processes are also seen around the facial nerve (7N)(arrowhead). Processes in the medial facial nerve area do not colocalize with VACHT antibody (J’’). Brain pictures taken from Mouse Brain Atlas (Paxinos and Franklin, 2007). Abbrev: SPN, sympathetic preganglionic nucleus; 7N, facial nerve; Int, interpositus; IRt, intermediate reticular nucleus; Sp5, spinal trigeminal nucleus; Amb, ambiguus nucleus; PFl, paraflocculus; PML, paramedian lobule. Scale bars: 100 μm unless otherwise noted; D, E, F’, and J’’ insets are 10 μm; F and I are 1 mm.

The first cross used the *Hoxa4::Cre* and *Chat*^*IRES-FLPo*^ alleles crossed to the intersectional tdTomato reporter (*R26*^*LSL-FSF-tdTomato*^, Ai65)(hereafter called *Chat*^*Hoxa4*^ neurons)(Madisen et al., 2015). *Hoxa4::Cre* is expressed in regions caudal to the developing rhombomeres 6/7, which corresponds to the caudal medulla (Huang et al., 2012; Yuengert et al., 2015), and the *Chat*^*IRES-*FLPo^, generated by Gord Fishell and Rob Machold (unpublished), is expressed in cholinergic neurons. Therefore, this cross should label all cholinergic neurons caudal to the lower medulla including spinal cord. We found that approximately 12.5% (mean, range 10-15%, n=2) of CHAT^+^ MN pools were labeled in this cross (Fig. 5B, representative images). When we analyzed the hindbrain sections of two samples, we found consistent axonal labeling in the intermediate reticular nucleus (IRt), facial nucleus (7N), spinal trigeminal nucleus (Sp5), and ambiguus nucleus (Amb)(Fig. 5C, orange and green triangles). The processes in the IRt were cholinergic as the TOM^+^ processes colocalized with the vesicular acetylcholine transporter (VACHT) antibody (Fig. 5F-F’, arrows). For Sample 1, we found TOM^+^ processes in the vermis of folia III and paraflocculus (PFl). Curiously, these processes did not express excitatory marker vesicular glutamate transporter 2, VGLUT2 (VG2), and resided between the NEUN^+^ granule cells (Fig. 5D, E). We were unable to assess whether these processes were cholinergic because we did not get reliable VACHT antibody staining in the granule cell layer of the cerebellum. Other areas of the hindbrain that had TOM^+^ processes are annotated for both samples in Fig. 5C. We note that axon terminations of sensory neurons labeled in the superficial dorsal horn of *Chat*^*Hoxa4*^ mice can be seen (Fig. 5B, arrowheads) and are likely due to transient expression of *Chat*^*IRES-*FLPo^ because *Chat* is not highly expressed in sensory neurons (Sharma et al., 2020).

To obtain a three-dimensional view of the axonal trajectories, we cleared the spinal cord and brain of a *Chat*^*Hoxa4*^ mouse using SHIELD (Park et al., 2018) and imaged with light sheet microscopy. In this cleared sample, MNs and other cholinergic neurons in the spinal cord such as the sympathetic pre-ganglionic nucleus (SPN) and V_0c_ neurons are labeled (Zagoraiou et al., 2009; Deuchars and Lall, 2015)(Movies 2 and 3). When examining the hindbrain for axonal projections, we found almost no labeling except for some asymmetric TOM^+^ labeling in Crus I and Crus II on only one side of the cerebellum of unclear origin (Fig. 5C, purple triangles, Movie 1). The cleared *Chat*^*Hoxa4*^ sample had cell bodies labeled in what appears to be the accessory facial nerve (acs7) and facial nerve (7N)(Movie 1) and had some processes in Sp5. In the spinal cord of the *Chat*^*Hoxa4*^ cleared sample, we found most MNs had axonal projections heading to the ventral root (Movies 2 and 3). However, occasionally, axons heading rostrally were seen, particularly in the cervical-thoracic cleared spinal cord (Movie 2) where one prominently labeled axon was seen on the left side of the spinal cord (right side of the image). When following the fluorescence of this axon, it appeared to have originated from the contralateral side of the spinal cord, but the fluorescence disappeared making it impossible to attribute this axon to any given neuron.

Due to the sparse labeling of MNs in the *Chat*^*Hoxa4*^ mice, it was difficult to conclude whether a lack of consistent axonal projections in the cerebellum was due to a lack of ascending projections or lack of sufficient labeling. Therefore, we pursued a second intersectional cross using *Chat*^*IRES-Cre*^ and *Cdx2::FLPo* crossed to the intersectional Ai65 tdTomato reporter (hereafter called *Chat*^*Cdx2*^ neurons). *Chat*^*IRES-Cre/+*^ labels all cholinergic cells and cells with transient *Chat* expression (Rossi et al., 2011; Nasirova et al., 2020) while the *Cdx2::FLPo* labels all cells caudal to the mid-cervical area of the spinal cord (Bourane et al., 2015). We found that in the spinal cord, labeling of MNs was much more robust with 92% of the MNs being labeled (n=1) (Fig. 5H). Similar to the *Chat*^*Hoxa4*^ mice, we saw axonal projections from sensory neurons in the dorsal horn indicating some transient expression of *Chat*^*IRES-Cre*^ in sensory neurons (Fig. 5H, arrowhead). In this mouse, we found many mossy fiber-like terminations in vermis II and III (Fig. 5I-I’, J-J’). In addition, we found prominent TOM^+^ processes in the medial part of the facial nerve (7N)(Fig. 5J’’ and insets, arrowhead) that were not cholinergic (VACHT^-^).

We note here that we pursued a third intersectional cross of *Atoh1*^*Cre/+*^ and *Chat*^*IRES-FLPo*^ alleles (unpublished). This cross labeled known *Atoh1*-lineage cholinergic neurons in the pedunculopontine tegmentum (PPTg) and the lateral dorsal tegmentum LDTg)(unpublished and Rose et al., 2009). However, PPTg neurons have been shown to project to deep cerebellar nuclei (Woolf and Butcher, 1989; Jaarsma et al., 1997), thus, confounding any interpretation of potential *Atoh1*-lineage cholinergic projections from the spinal cord.

### Infection of Purkinje cells by injection of viruses in the periphery

As an alternative strategy to test whether IF MNs could send ascending axon collaterals, we sought to isolate specifically the IF MN pool and trace their arborizations. We reasoned that injection of CRE viruses into the hindpaw of tdTomato (TOM) reporter mice (*R26*^*LSL-tdTom*^, Ai14)(Madisen et al., 2010) would infect the MN axons allowing for TOM expression in the entire MN. If any ascending axon collaterals existed, we would see TOM^+^ labeling in the cerebellum. We performed multiple experiments using both AAV and Lentivirus (AAV8-GFP-Cre and Lenti^FugE^-Cre) at early (P4-5) and later (P14-15) time points with 21-27 days allowed for expression (Fig. 6). We chose to use AAV8, which was reported to have sparse infection of the MNs at P1 (Foust et al., 2008). Sparse labeling of MNs would allow us to trace axonal trajectories. In addition, although AAVs have been reported to cross the blood-brain barrier (BBB) at early postnatal stages, those experiments were performed with intraperitoneal or intravenous infection, whereas we were targeting specifically the hindpaw area as well as later time points (P14-15) (Foust et al., 2008; Foust et al., 2009; Gray et al., 2011; Zhang et al., 2011). Furthermore, we tested infection with Lentivirus, which to the best of our knowledge, is not known to cross the BBB.

**Figure 6.**
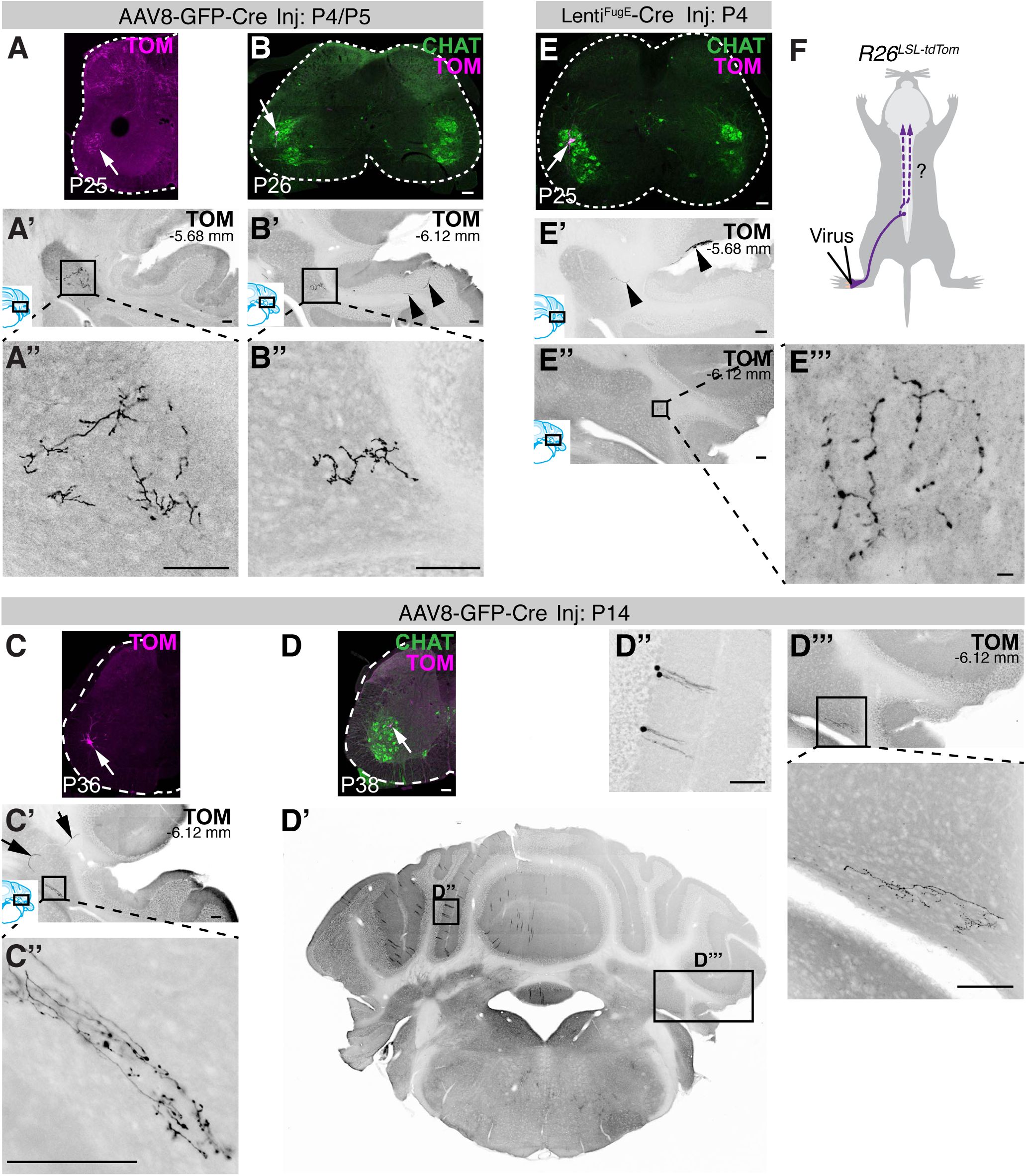
Peripheral injections of viruses infect Purkinje cells in the cerebellum. (A-B’’) Injection of AAV8-GFP-Cre at P4 or P5 into the hindpaw of two representative animals (*R26*^*LSL-tdTom*^) and harvested 21 days later shows TOM fluorescence within the IF MN pool indicating infection and recombination (A, B, arrows). Cerebellar sections show axonal processes in the dentate nucleus (A’-B’’). Labeling of a long-projecting axon can be seen (B’, arrowheads). (C-D’’’) Injection of AAV8-GFP-Cre at P14 into the hindpaw of two representative animals (*R26*^*LSL-tdTom*^) and harvested 22-24 days later shows TOM fluorescence within the IF MN pool indicating infection and recombination (C, D, arrows). Long-projecting axons are seen terminating in the dentate nucleus (C’-C’’, D’’’). In one example injection, Purkinje cells were infected on the same side as injection (D’, D’’) and axons projecting to the dentate were seen on the contralateral side. (E-E’’’) Injection of Lenti^FugE^-Cre into the left hindpaw of *R26*^*LSL-tdTom*^ mice at P4 and harvested 21 days later showed infection of the IF MN Pool (E, arrow). A Purkinje cell can be seen extending an axon to the dentate nucleus (E’, arrowheads) and its terminals in the dentate nucleus seen in a more caudal section (E’’’). (F) Schematic of the proposed hindpaw injections. MN pools identified with CHAT Antibody in B, D, and E. Brain pictures taken from Mouse Brain Atlas (Paxinos and Franklin, 2007). Distance from Bregma given in mm. Scale bars: 100 μm, E’’’ is 10 μm.

In all the injections, consistent with infection of MN axons innervating the hindpaw, we found a small number of TOM^+^ neurons in the IF MN pool (Fig. 6A,B, C, D, E, arrows). In the cerebellum, we found that Purkinje cells expressed tdTomato upon unilateral injection of viruses into the hindpaw. In all cases, axonal arborizations in the dentate nucleus could be seen (Fig. 6A’’, B’’, C’’, D’’’, E’’’). In some cases axons could be seen traveling to the dentate (Fig. 6B’, C’, E’, arrowheads) and a Purkinje cell (PC) cell body was labeled (Fig. 6E’, arrowhead). In general, PC labeling was quite sparse except in one case where many PCs were labeled on the same side as the injected hindpaw and the axons projecting to the dentate were on the contralateral side to the injected hindpaw (Fig. 6D’-D’’’). In the two cases where we preserved the orientation of the cerebellum relative to the injection site, we found that the axons in the dentate were on the side contralateral to the hindpaw injection site (Fig. 6D’’’, E’’’). Altogether, using viral tracing strategies, we could label the IF MNs; however, axons seen in the cerebellum appeared to come from infected PCs on the periphery of the cerebellum.

### Retrograde labeling from the cerebellum

Lastly, to test whether IF MNs could send axons to the cerebellum, we injected the retrograde tracer Fluorogold (FG) into the vermis of folia I-V in the cerebellum (Fig. 7A-B). In the IF MN pool of a FG injected mouse, none of the *Atoh1*-lineage MNs (TOM^+^CHAT^+^) were co-labeled with FG (arrowheads)(Fig. 7C). Notably, other FG labeled cells in the ventral spinal cord, some of which have a large cell body morphology like MNs, are not cholinergic (FG^+^CHAT^-^)(Fig. 7D’’’-D’’’’, arrowheads). The FG-retrograde labeling worked successfully, though, as other cerebellar-projecting neurons in the medulla and spinal cord, namely, the external cuneate nucleus (ECu), lateral reticular nucleus (LRt), inferior olive (IO), and Clarke’s column (CC)(Fig. 7B, D) were labeled. Furthermore, consistent with our previous findings (Yuengert et al., 2015), an occasional CC cell is *Atoh1*-lineage (FG^+^TOM^+^, Fig. 7D’, arrow). Interestingly, only one of the cells in the cluster of retrogradely-labeled cells lateral to CC is *Atoh1*-lineage (FG^+^TOM^+^, Fig. 7D’’, arrow) indicating that few of the *Atoh1*-lineage neurons target Folia I-V in the cerebellar vermis. Two female mice were injected with FG with similar results, so a representative animal (mouse #2) is shown in Fig. 7.

**Figure 7.**
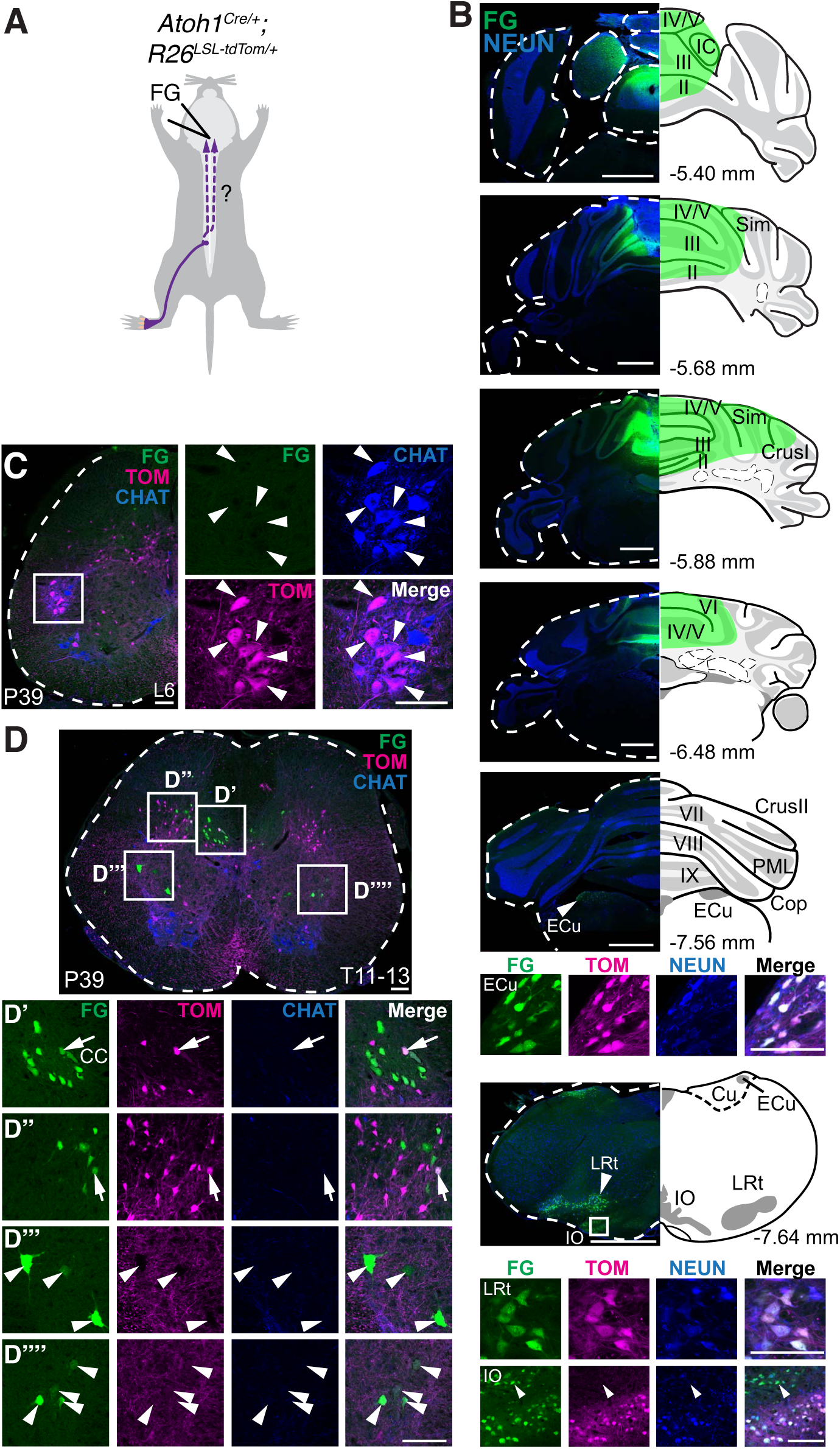
Injection of Fluorogold retrograde tracer into the cerebellar vermis does not label the IF MN pool. (A) Schematic of Fluorogold (FG) injections into the cerebellar vermis of *Atoh1*^*Cre/+*^ knock-in mice crossed to the TOM reporter mice. (B) FG (green) was injected into the vermis of folia II-V. The injections were symmetrically spread from the midline, so only one half is shown. Areas in the medulla that are known to project to the cerebellum (ECu, LRt, and IO) are retrogradely labeled (FG^+^). Many of the neurons in the ECu, LRt, and a portion of the IO (insets) are also *Atoh1*-lineage (TOM^+^) as previously reported (Rose et al., 2009). Note that a portion of the IO (arrowhead in bottom IO panel) are not TOM^+^. (C) There are no FG^+^ cells in the IF MN pool (FG^-^TOM^+^CHAT^+^, arrowheads). (D-D’’’’) Cerebellar-projecting CC cells and cerebellar-projecting cells lateral to CC are retrogradely labeled with FG (D’-D’’). Few of these retrogradely labeled cells are *Atoh1*-lineage (FG^+^TOM^+^, arrows in D-D’’). Cerebellar-projecting cells in the ventral spinal cord are not *Atoh1*-lineage and are not cholinergic (D’’’-D’’’’, FG^+^TOM^-^CHAT^-^, arrowheads). Brain pictures taken from Mouse Brain Atlas (Paxinos and Franklin, 2007). Abbrev: IC, inferior colliculus; Sim, Simplex; PML, paramedian lobule; ECu, external cuneate nucleus; LRt, lateral reticular nucleus; IO, inferior olive; Cu, cuneate; CC, Clarke’s column. Distance from Bregma given in mm. Scale bars: 100 μm in all panels except cerebellar sections in B are 1 mm.

## Discussion

The impetus for this study stemmed from the serendipitous finding that IH and IF MNs are labeled in *Atoh1*^*Cre/+*^ mice. This finding led to two lines of inquiry: the development as well as the potential novel connectivity of IH and IF MNs.

### IH and IF MNs have a unique developmental history

We found that *Atoh1*^*Cre/+*^ mice label the IH and IF MNs consistent with these MN pools having a unique developmental history. Other MN pools such as the gastrocnemius and the tibialis anterior are not enriched in the *Atoh1*^*Cre/+*^ mouse line indicating that the IH and IF MN pools have distinct molecular processes that allow for tdTomato expression. We were able to isolate labeling of the IH and IF MNs in the *Atoh1*^*Cre/+*^ mouse line to the postnatal time period, although the levels of *Atoh1* mRNA at the times tested were below detectable limits. However, the consistent and robust labeling of IH and IF MNs in the *Atoh1*^*Cre/+*^ mouse line indicates that these MNs have a unique developmental program compared to the neighboring LMC MN pools that are not labeled.

Although *Atoh1*^*Cre/+*^ is enriched in the IH and IF MN pools, the cell type of the MNs labeled by the *Atoh1*^*Cre/+*^ mouse line is heterogeneous. We found that *Atoh1*^*Cre/+*^ TOM^+^ IH and IF MN pools represent a mixture of α- and γ-MNs and with only a subset expressing a marker for fast twitch MNs. The *Atoh1*^*Cre/+*^ TOM^+^ axons innervate both the intrafusal and extrafusal muscle of the hindpaw lumbrical muscle consistent with both α- and γ-MNs being labeled. It is also possible, though, that *Atoh1*^*Cre/+*^ TOM^+^ MNs are labeling β-MNs. β-MNs are MNs whose axons bifurcate to simultaneously innervate both intrafusal and extrafusal muscle (Manuel and Zytnicki, 2011). Currently, to the best of our knowledge, there are no known molecular markers for β-MNs that would allow us to test whether the *Atoh1*^*Cre/+*^ TOM^+^ MNs are β-MNs. Interestingly, one of the earliest descriptions of β-MNs in mammalian systems was from the lumbrical muscle of cats (Bessou et al., 1965) and could be a feature of IH and IF MNs. It is enticing to speculate that the MNs involved in fine motor skill would have β-MNs as a feature because functionally, β-MNs would allow for simultaneous contraction of intrafusal and extrafusal muscle, thus resetting the muscle spindle to respond immediately to stretch activation (Manuel and Zytnicki, 2011).

### MNs as a source of corollary discharge

The finding that *Atoh1*, a transcription factor that specifies many cerebellar-projecting neurons, could be enriched in the IH and IF MNs led to the intriguing hypothesis that *Atoh1* might act in these MNs to send axon collaterals to the cerebellum. A model by which MNs themselves could send a copy of their motor information directly back to the cerebellum or other hindbrain areas was enticing. Such a model would suggest that motor information, in particular fine motor information, would need to be immediately sent back to the cerebellum as a means of updating the proprioceptive sensory system about self-generated movements. Furthermore, having a corollary discharge pathway specifically from digit-innervating MNs might make temporal sense given that fine motor movements would need more rapid updating of the sensory-motor system compared to whole limb movements. We attempted to test the hypothesis that IH and IF MNs could send ascending axon collaterals using intersectional genetics, viral labeling, and retrograde labeling strategies, but were unable to find evidence to support this idea.

### Intersectional genetic labeling of caudal cholinergic neurons

To label caudal cholinergic neurons, we pursued two intersectional strategies generating *Chat*^*Hoxa4*^ and *Chat*^*Cdx2*^ mice, both of which had limitations. The labeling of MNs in the *Chat*^*Hoxa4*^ mice was too sparse (12.5%) to ensure that the IH and IF MN pools were sufficiently labeled. The lack of robust MN labeling in the *Chat*^*Hoxa4*^ mice might be due to the dorsal-enriched expression of *Hoxa4::Cre* that would miss ventrally generated neuronal populations (Yuengert et al., 2015). In contrast, the *Chat*^*Cdx2*^ mice labeled almost all the MNs (92%), but also labeled many other neurons with potentially ascending projections in the spinal cord likely due to transient *Chat* expression (Nasirova et al., 2020). In both mice, other cholinergic neurons in the spinal cord (SPN, V_0c_) were labeled, although these should not have any ascending projections (Zagoraiou et al., 2009; Deuchars and Lall, 2015).

Taking both *Chat*^*Hoxa4*^ and *Chat*^*Cdx2*^ mice together, the only places that had common hindbrain projections were in folia III of the cerebellar vermis and in the facial nerve nucleus (7N). The projections to the facial nucleus were not cholinergic (Fig. 5J’’) and are likely evidence of transient cholinergic expression in these neurons that become glutamatergic in adulthood (Nasirova et al., 2020). The projections to folia III are of unknown neurotransmitter status as antibodies for VACHT do not work well in the cerebellar granule cell layer and these TOM^+^ projections in *Chat*^*Hoxa4*^ mice did not express VGLUT2. Therefore, it is difficult to identify the origin of the projections to folia III and it is possible that they also come from cells that transiently express *Chat*. Note the many TOM^+^ cells in the spinal cord that do not colocalize with CHAT antibody (Fig. 5H) representing transiently expressing *Chat* neurons.

As for other ascending projections in either the *Chat*^*Cdx2*^ or *Chat*^*Hoxa4*^ mouse, the TOM^+^ processes in IRt are not as prominent in the *Chat*^*Cdx2*^ mouse as in the *Chat*^*Hoxa4*^ mouse indicating that these cholinergic IRt processes in the *Chat*^*Hoxa4*^ mice might be from cells labeled between the caudal medulla and mid-cervical spinal cord. This would suggest that the cholinergic innervation of IRt neurons involved in respiration might come from the cervical spinal cord area (Anderson et al., 2016; Nasirova et al., 2020). In the *Chat*^*Hoxa4*^ mice, we found areas of the medulla that were previously reported to contain cholinergic projections in *Chat*^*IRES-Cre/+*^ mice (facial nucleus (7N), spinal trigeminal nucleus (Sp5), ambiguus nucleus (Amb), and paraflocculus (PFl))(Nasirova et al., 2020) and were able to isolate these projections as originating from cell bodies caudal to the lower medulla. Furthermore, to get a three-dimensional view of MN axonal trajectories that could be missed in thin cryosections, we cleared the spinal cord and hindbrain of a *Chat*^*Hoxa4*^ mouse. In the *Chat*^*Hoxa4*^ mouse, we saw one ascending projection in the spinal cord and some faint asymmetric TOM^+^ labeling in right Crus I and II, but could not follow the axons of these to identify the originating cell. Altogether, even though we saw some projections in cerebellar vermis III in both, *Chat*^*Hoxa4*^ and *Chat*^*Cdx2*^ mice, it remains ambiguous as to whether these projections originated from MNs due to the large number of transiently-expressing *Chat* neurons that were labeled.

### Viral injections in the periphery infect Purkinje cells in the CNS

To analyze the projections specifically of IF MNs, we injected CRE AAV and Lentiviruses into the hindpaw of CRE-dependent TOM reporter mice. While this resulted in sparse labeling of the IF MN pool, when we analyzed the cerebellum, we found that Purkinje cells (PCs) were also labeled. The viruses were injected into the hindpaw where they presumably infected muscle, nerve, and blood vessels. For two animals where the orientation was recorded, the PCs contralateral to the injected hindpaw were infected.

There are at least two potential explanations for how peripheral injection of viruses infect PCs in the central nervous system (CNS). One possible explanation is that CRE-expressing AAVs are “hopping” one synapse in an anterograde fashion as has been previously reported (Zingg et al., 2017). In this scenario, the CRE AAV could potentially reflect a connection between the initially infected MN and PCs in the cerebellum. We think this possibility is unlikely given that one mouse had a PC labeled on the edge of Crus I (Fig. 6E’) and another on the edge of paraflocculus (Fig. 6B’) indicating that the infection is random. In addition, we saw the same infection of PCs using a CRE-expressing lentivirus, which has not been reported to travel transynaptically. A second possible explanation is that the viruses can travel across the blood brain barrier (BBB) to preferentially infect PCs in the CNS. This would explain why several PCs were labeled in one mouse on both the contralateral and ipsilateral sides and yet we do not see widespread infection of other cell types (Fig. 6D’).

Several AAV serotypes including AAV8 have been reported to cross the BBB, particularly at early postnatal stages (P1) when the BBB is not yet fully developed (Foust et al., 2008; Foust et al., 2009; Gray et al., 2011; Zhang et al., 2011). AAV8 was reported to sparsely label MNs, which we thought would work to our advantage in tracing single neuronal axons (Foust et al., 2008). In addition, we reasoned that local delivery to the hindpaw, rather than the more global intraperitoneal and intravenous injections described previously, would limit crossing the BBB. However, we found that infection of the CNS using AAV8 and Lentivirus at postnatal stages slightly later than P1 (namely P4-5 and P14-15) can cross the BBB sparsely infecting PCs. We present the data here as a warning for future studies looking a peripheral to CNS connectivity.

### Reconciling spinocerebellar retrograde tracing studies

Several retrograde-labeling studies of spinocerebellar neuron in cats, rats, opossum, and monkeys have identified cerebellar-projecting cells in spinal cord lamina IX where MNs reside (Cooper and Sherrington, 1940; Matsushita and Hosoya, 1979; Matsushita et al., 1979; Terman et al., 1998). Together with our findings that *Atoh1*-lineage neurons labeled IF MNs, we tested the hypothesis that IF MNs could send ascending projections to the cerebellum. However, our injections of FG into the cerebellar vermis did not label the IF MN pool (Fig. 7). Furthermore, cerebellar-projecting cells in the ventral spinal cord, some of which had a large cell body morphology similar to MNs, were not cholinergic. Therefore, the developmental origin of these ventral cerebellar-projecting cells remains unknown. Although evidence for IF MNs sending axon collaterals to the cerebellum is not supported by our data, it is possible that potential ascending projections terminate in other areas of the cerebellum or hindbrain that were not targeted in our FG injections.

Altogether, we present here that the *Atoh1*^*Cre/+*^ mouse consistently labels MNs of the IH and IF and that the *Atoh1*^*Cre/+*^ mouse could be used to specifically label these populations. Whether the IH and IF MNs send axon collaterals to the cerebellum as a mechanism of corollary discharge, as hypothesized, remains an open question. We tested this hypothesis using three distinct approaches: intersectional genetics, viral labeling, and retrograde labeling, but were unable to find evidence to support the idea that IH and IF MNs send axon collaterals to the cerebellum.

## Acknowledgments

This work was supported by P01 NS074972 to G.F. and R.P.M; R21 NS099808, R01 NS100741, and start up funds to H.C.L. We thank Sam Pfaff for the HB9 antibody, Tom Jessell for the Cpne4 and Fign antibodies, Lin Gan for the *Atoh1*^*Cre/+*^ knock-in mouse, Martyn Goulding for the *Cdx2::FLPo* mouse, Thomas Südhof for the Syntaxin1 antibody, and Heankel Cantu Oliveros and Wei Xu for the Lenti^FugE^-Cre virus, the Neuroscience Microscopy Facility which is supported by the UTSW Neuroscience Dept. and the UTSW Peter O’Donnell, Jr. Brain Institute, Katherine Casey, Clayton Trevino, Megan Goyal, and Adelaide Brooks for technical assistance, LifeCanvas Technologies for tissue clearing assistance, Kevin Dean for computer use, Peter Tsai, Todd Roberts, and Julian Meeks for helpful discussions, and Weichun Lin, Jon Terman, and Jane Johnson for careful reading of the manuscript.

## Multimedia, Figure, and Table

**Movie 1.** Movie of the tdTomato labeled cells in the hindbrain of a SHIELD cleared *Chat*^*Hoxa4*^ mouse. Some punctate fluorescence is seen in right Crus I.

**Movie 2.** Movie of the tdTomato labeled cells in the cervical to thoracic spinal cord of a SHIELD cleared *Chat*^*Hoxa4*^ mouse. Most MN axons appear to go into the ventral roots. One axon on the left side of the spinal cord (right side of the image) appears to originate from the contralateral side of the spinal cord, but the fluorescence cannot be traced to the original neuron.

**Movie 3.** Movie of the tdTomato labeled cells in the thoracic to lumbar spinal cord of a SHIELD cleared *Chat*^*Hoxa4*^ mouse. Most MN axons travel to the ventral roots.

